# Dynamic Regulation of RNA Structure in Mammalian Cells

**DOI:** 10.1101/399386

**Authors:** Lei Sun, Furqan Fazal, Pan Li, James P. Broughton, Byron Lee, Lei Tang, Wenze Huang, Howard Y. Chang, Qiangfeng Cliff Zhang

**Affiliations:** MOE Key Laboratory of Bioinformatics, Center for Synthetic and Systems Biology, Beijing Advanced Innovation Center for Structural Biology, Tsinghua-Peking Joint Center for Life Sciences, School of Life Sciences, Tsinghua University, Beijing 100084, China; Center for Personal Dynamic Regulomes, Stanford University, Stanford, CA 94305, USA; Howard Hughes Medical Institute

## Abstract

RNA structure is intimately connected to each step of gene expression. Recent advances have enabled transcriptome-wide maps of RNA secondary structure, termed RNA structuromes. However, previous whole-cell analyses lacked the resolution to unravel the dynamic regulation of RNA structure across subcellular states. Here we reveal the RNA structuromes in three compartments — chromatin, nucleoplasm and cytoplasm. The cytotopic structuromes substantially expand RNA structural information, and enable detailed investigation of the central role of RNA structure in linking transcription, translation, and RNA decay. Through comparative structure analysis, we develop a resource to visualize the interplay of RNA-protein interactions, RNA chemical modifications, and RNA structure, and predict both direct and indirect reader proteins of RNA modifications. We validate the novel role of the RNA binding protein LIN28A as an N6-methyladenosine (m^6^A) modification “anti-reader”. Our results highlight the dynamic nature of RNA structures and its functional significance in gene regulation.

---

RNAs fold into complex structures that are crucial for their functions and regulations including transcription, processing, localization, translation and decay^1-6^. Over the last few decades RNA structure has been studied extensively *in vitro* and *in silico*, and crystallography and cryo-EM structures of molecular machines such as the spliceosome and ribosome, containing RNAs at their core, have become available^7,8^. In recent years technologies have been developed to map RNA secondary structures for the whole transcriptome, *i.e*., RNA structuromes, by combining biochemical probing with deep sequencing^9-16^. These systems biology studies have revealed many novel insights on the RNA structure basis of gene regulation^17-20^. However, so far existing genome-wide structure probing studies have focused on whole-cell data, which only represents an ensemble average of RNA molecules in different subcellular compartments.

In fact, RNA undergoes a complex life cycle in eukaryotic cells, mirrored by its movement into distinct cytotopic locales^4,21^. RNA structure is thought to form co-transcriptionally on the chromatin template, undergo conformational changes resulting from RNA chemical modification and processing in the nucleus, and experience further changes in the cytoplasm during translation and RNA decay. Averaging the RNA structure signal in the entire cell may obscures these critical dynamic features. More importantly, detailed mapping of RNA structures *in situ* will help to elucidate how they are regulated, which is essential to understanding the RNA structure basis for gene expression regulation.

An important driving force that regulates the dynamic landscape of RNA structures in post-transcription regulation is the binding of RNA-binding proteins (RBPs). A study in Arabidopsis revealed that RNA secondary structure is anti-correlated with protein-binding density^22^. We recently used icSHAPE to probe RNA structuromes in mouse ES cells and examined the *in vivo* and *in vitro* structure profiles of RBFOX2, a splicing factor of the “feminizing locus on X” (Fox) family proteins; and HuR, an RBP that regulates transcript stability^12^. We implemented a machine learning algorithm and found that using structure signals significantly improved the prediction of RNA-binding sites of both RBPs, suggesting that RNA structure signature analysis is a powerful tool to investigate RNA–RBP interactions. However, in spite of these recent advances in our understanding of the association between RNA structure and RBP-binding, a compendium of the RNA structural basis of RBP binding is not available.

In addition to RBP binding, the modification and editing of RNAs are also an important mechanism for RNA structure regulation. RNA modification can regulate almost all RNA processes including RNA maturation, nuclear retention and exportation, translation, decay, and cell differentiation and reprogramming as well^23,24^. As one of the most abundant and important types of mRNA modification, N6-methyladenosine (m^6^A) has been showed to be able to weaken duplex RNA stability by conformational switching^12,25,26^. The impact of structure destabilizing effect of m^6^A is exemplified by a study that investigated HNRNPC, a splicing factor that preferentially binds to single-stranded polyU tracts^27^. Biochemical studies showed that m^6^A modification can disrupt the local RNA structures and promote HNRNPC binding in nearby regions^28^. The study defined these m^6^A sites as “m^6^A-switches”, and identified an enrichment of tens of thousands of m^6^A-switches in the vicinity of HNRNPC binding sites. The authors further showed these m^6^A-switches could regulate HNRNPC-binding and the splicing of the target RNAs. This study demonstrated a very nice example that m^6^A functions as an RNA structure modulator to affect RNA splicing through interfering with protein binding. However, whether the structure context is a generic mechanism for the recognition of other “reader” proteins of m^6^A and other RNA modifications, is still unclear^29^.

Here we use *in vivo* click selective 2-hydroxyl acylation and profiling experiments (icSHAPE)^12^, a technique we developed to map RNA structure *in vivo*, in three compartments – chromatin, nucleoplasm and cytoplasm – in both mouse and human cells. Consequently, we were able to determine the precise relationship of RNA structure with cellular processes including transcription, translation and RNA decay in the compartment where they occur. Separately, we could quantify how RNA adopts different conformations across different cellular compartments, which we termed “structural dynamics”, and investigate the sophisticated interplay of RNA structured dynamics, RNA modification and RBP binding.

## Results

### Cytotopic RNA structure maps substantially expand the scope and comprehensiveness of RNA structures

To investigate the dynamic regulation of RNA structure in the cell, we performed icSHAPE to measure RNA secondary structure for transcripts isolated from three subcellular compartments and in two species (Figure 1a). After performing the icSHAPE reaction of living cells *in situ*, RNA fractionation^30,31^ enabled the study of RNA structural changes in distinct subcellular locations. Separately, we fractionated the three subcellular compartments, isolated and refolded naked RNA from each, and performed icSHAPE *in vitro*. This *in vitro* dataset served as a control for the RNA contents in each compartment. The use of both v6.5 mouse embryonic stem (mES) cells and human embryonic kidney (HEK293) cells allowed us to examine whether the structural patterns we observed are conserved across the two species and cell types.

**Fig 1.**
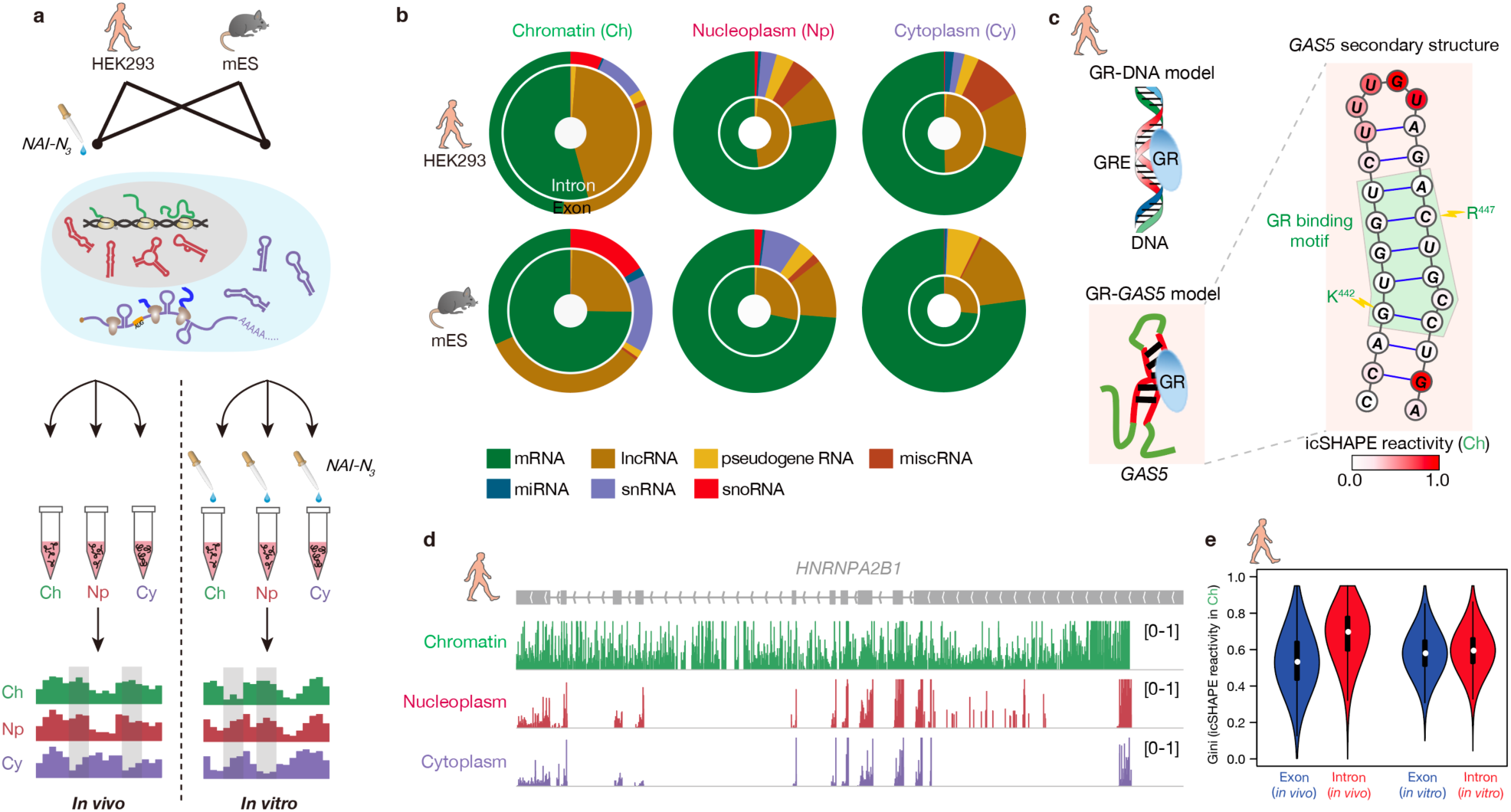
Chromatin fractions are enriched for pre-mRNA and lncRNA structures. **a**, Experimental overview of the icSHAPE protocol. **b**, Donut charts showing read distributions of different RNA types in the three cellular compartments. The outer circles represent exon coverage while the inner ones represent intron coverage. **c**, *GAS5* RNA secondary structure with icSHAPE reactivity scores shown in color. The nucleotides outlined in red interact with the GR amino acids, as shown in blue. **d**, UCSC tracks showing icSHAPE reactivity scores (y-axis), along the RNA sequence. Here 1 denotes unstructured (single-stranded) regions, and 0 denotes fully-structured regions. **e**, Violin plot of Gini index of icSHAPE data in exon versus in intron.

We determined RNA structure, as previously described^12,32^, after enriching for messenger RNAs (mRNAs) and long noncoding RNAs (lncRNAs) by ribosome depletion, and sequencing the resulting icSHAPE libraries at high depth (∼200 million reads per replicate, Table S1). We first confirmed the quality of fractionation using RT-qPCR for landmark RNAs, and Western blots for specific proteins (Figure S1). We used the icSHAPE pipeline^12^ to calculate a score to represent the structural flexibility (indicative of unpaired RNA bases) of every nucleotide, and found good correlation across replicates (Pearson correlation *r* > 0.75 for the top 60% most-abundant transcripts in all replicates, Figure S2). To further validate our structural data, we examined its agreement with known structures – two such RNAs are *Ribonuclease P* RNA and Signal recognition particle (*SRP*) RNA (Figure S3). Both RNAs are enriched in nucleoplasm, and indeed our nucleoplasm icSHAPE data closely match the existing structural models.

The chromatin structurome is enriched for lncRNAs (Figure 1b, S4a). As an example, we examined the structure of the human growth arrest-specific 5 (*GAS5*) noncoding RNA, which acts as a decoy glucocorticoid response element (GRE) by binding to the DNA-binding domain of the glucocorticoid receptor (GR)^33^. Indeed, the expected *GAS5* RNA structure is accurately recovered in the chromatin fraction, showing low icSHAPE scores for the double-stranded GR binding motif of the *GAS5* RNA, and high reactivity score for the loop region (Figure 1c). Similar to lncRNAs, snoRNAs and snRNAs are also enriched in the chromatin fraction, and to a smaller extent in the nucleoplasm fraction (both relative to the cytoplasm). Furthermore, intronic reads constitute the majority of the sequencing data in the chromatin fraction, but only ∼15-20% of reads in the cytoplasmic fraction (Figure 1b, Figure S4b). For example, we obtained intron structures for the transcript heterogeneous nuclear ribonucleoprotein A2/B1 (*HNRNPA2B1*) in the chromatin fraction, but these sequences were largely absent in the nucleoplasmic and cytoplasmic fractions (Figure 1d). Interestingly, we found that RNAs *in vivo* are much more folded in intron regions than in exon regions (average Gini index of 0.7 versus 0.5. A higher Gini index indicates a more structured region^11^), in contrast to *in vitro* conditions (both with average Gini index 0.6); this result holds true for both human (Figure 1e) and in mouse (Figure S5a). The finding that intronic regions are more folded *in vivo* is not likely due to differential RNA-binding-protein (RBP) binding in introns versus exons, as similar trends were observed when all known RBP-binding sites were excluded in the structural comparison (Figure S5b). Instead, these results may suggest distinct interplays between RNA structures, and transcriptional or splicing regulation in introns and exons. In summary, the RNA-structural profiles of the chromatin fraction provide a rich resource to interrogate structures of lncRNAs, pre-mRNAs including introns, and other chromatin-associated RNAs, expanding the scope of the RNA structurome.

### RNA structure plays a central role in connecting many cellular events

The cytotopic RNA structuromes allowed us to assess the roles of RNA structure (or lack thereof) in regulating each step of the gene-expression life cycle, which takes place in distinct subcellular compartments. We obtained data on transcriptional rate, translational efficiency, and RNA half-life from previous studies in human and mouse^34-36^, and correlated data with the Gini index of icSHAPE reactivity. RNA structure in nascent RNA has been suggested to propel or impede RNA polymerase pausing at individual genes^37^. We therefore analyzed the relationship between transcription and 5’UTR (untranslated region) RNA structures of the chromatin-associated fraction, and found that lower transcriptional rate correlates modestly with more structure (*r* = – 0.19, *p* = 1.5 × 10^−6^, Figure 2a). Next, many studies have found RNA secondary structure upstream of or at ribosome binding sites is important for translation^15,17-19^. We did observe that more 5’UTR RNA structure correlates with decreased translational efficiency in the cytoplasmic fraction (*r* = –0.31, *p* = 1.7 × 10^−48^, Figure 2b). Finally, as RNA degradation occurs in both nucleoplasm and cytoplasm via different pathways, we analyzed the dependence of RNA half-life on RNA structure in both fractions. We found that more-structured RNAs tended to have shorter half-lives in both the nucleus and cytoplasm (Figure 2c-d), highlighting the emerging and under-appreciated role of double-strand ribonucleases in transcript turnover^38^.

**Fig 2.**
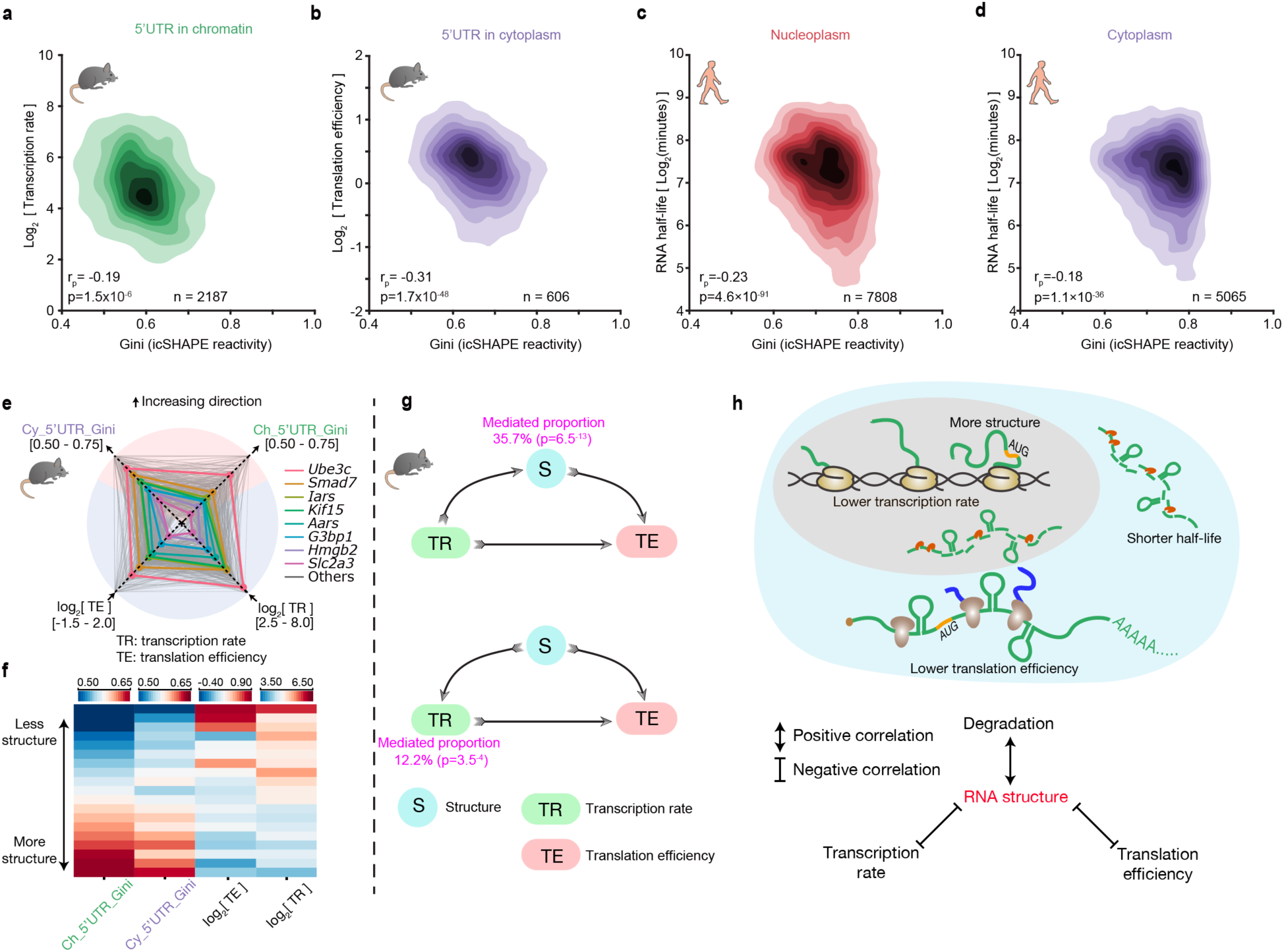
RNA structure plays a central role in connecting transcription, translation and RNA degradation. **a-d**, Scatter plots of (a) transcription rate versus 5’UTR RNA structure in chromatin, (b) translational efficiency versus 5’UTR RNA structure in cytoplasm, (c) RNA half-life versus full-length-transcript RNA structure in nucleoplasm, and (d) RNA half-life versus RNA structure in cytoplasm. The 2-tailed p-value was calculated by python package function *scipy.stats.pearsonr*. **e**, Radar diagram showing 5’UTR RNA structure in chromatin, 5’UTR RNA structure in cytoplasm, transcription rate, and translational efficiency. Grey lines show all genes, and the colored lines highlight representative transcripts. **f**, Heatmap of 5’UTR RNA structure in chromatin, 5’UTR RNA structure in cytoplasm, transcription rate, and translational efficiency. Each strip represents an average of a bin comprising 5% data, ranked by RNA-structure reactivity in the chromatin fraction, 477 common transcripts are showed here. **g**, Mediator model (above) and cofounding model (bottom) of RNA structure in connecting transcription rate and translation efficiency. P-values were calculated by double-ended t-test. **h**, Schematic showing RNA structure connects transcription, translation and RNA degradation.

Quantitative correlation analysis showed that the relationships among RNA structure, transcription and translation are not binary, as there is a general trend that an RNA with lower transcriptional rate tends to simultaneously be more structured and translated less efficiently (Figure 2e-f). The positive link between transcription and translation, two major events in gene expression, has been previously appreciated^39^(Figure S6). Recent studies have suggested different mechanisms, including m^6^A modification, that could account for this linkage by imprinting an mRNA transcript during its synthesis and later regulating its translation^39-41^. Our data suggest that genome-wide RNA structures formed at chromatin during transcription remain largely unchanged in the nucleoplasm and cytoplasm fractions (Figure S7), and might thus serve as a link between transcription and translation efficiencies. We therefore considered two models to explain our observations – in the first model RNA structure is a mediation factor that is affected by transcription, and it in turn affects translation; and in the second model RNA structure is a cofounding factor that has an effect on both transcription and translation (Figure 2g). Statistical analysis suggests that while both models could be true, the first (mediation) model can account for a larger fraction of the positive correlation between transcription and translation, and is statistically more significant. In summary, RNA structure plays a general role that connects many cellular events including transcription, translation and RNA degradation (Figure 2h).

### Pervasive RNA structure dynamics across different cellular compartments

More importantly, cytotopic RNA structuromes also enabled us to examine how RNA adopts different conformations across different cellular compartments, which we term “structural dynamics”. Overall, we saw that RNA structures were more unfolded in the chromatin fraction relative to the cytoplasmic fraction, even at sites not targeted by RBPs (Figure S8). As specific regions of an individual RNA can be regulated differently and display different patterns of structural dynamics, we implemented a statistical method to discover structurally-dynamic regions. As an example, we show that U12 snRNA displayed structurally-dynamic regions between compartments (Figure 3a, black bars). In addition, despite high evolutionary conservation of U12, the RNA structures showed shared and unique conformational changes in human and mouse. These findings suggest that both species-specific and conserved mechanisms may regulate RNA structures and structural dynamics.

**Fig 3.**
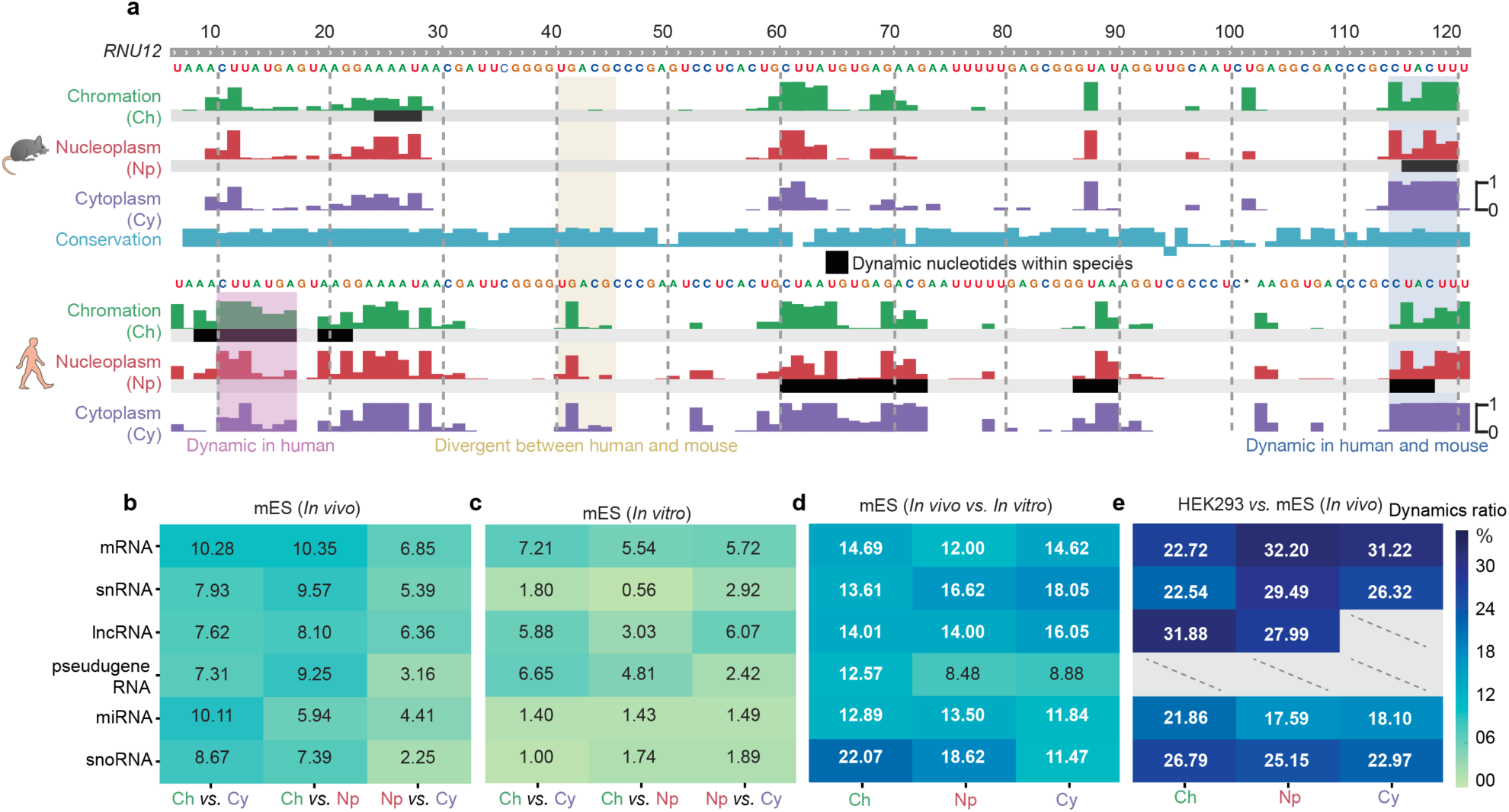
RNA structure differences in cellular context. **a**, U12 small nuclear RNA (snRNA) structure dynamics across cellular compartments, and the structural divergence in two species. Tracks show the icSHAPE score plotted along the RNA sequence. The black bars highlight RNA structurally-dynamic regions. **b-e**, Heatmaps showing fractions of structurally-different regions across cellular compartments (b) *in vivo*, (c) *in vitro*, (d) between *in vivo* and *in vitro*, and (e), between human and mouse. Dashed lines represent insufficient data.

On a genome-wide scale, we found that different RNA categories showed different levels of structural dynamics *in vivo* (Figure 3b). To begin to dissect the factors that regulate RNA structural dynamics in cells, we used the same analysis pipeline to evaluate data obtained from fractionated, purified RNA that was refolded *in vitro* (Figure 3c), and compared RNA conformational changes observed between compartments *in vivo* and *in vitro*. Conceptually, structural differences between RNAs isolated from different compartments and refolded in vitro do not represent structural dynamics, but we will use this term for simplicity. In general, as expected, RNA structures vary less between the compartments *in vitro* relative to *in vivo* (comparing Figure 3b to Figure 3c), suggesting that fewer factors influence RNA folding *in vitro* versus *in vivo*. This finding is particularly true for highly-conserved small RNAs such as snoRNAs, miRNAs and snRNAs, suggesting that these functional RNAs adopt stable structures *in vitro* but are subjected to extensive regulation *in vivo.* The structural differences are magnified when directly comparing *in vivo* to *in vitro* icSHAPE data for each compartment (Figure 3d), and different RNA categories displayed varying levels of structural differences *in vivo* and *in vitro*, consistent with previous findings from whole-cell data^12^. Finally, we compared the levels of structural divergence between mouse and human for sequence-conserved regions. We used the same pipeline used above to call for regions of structural changes, and found even larger fractions of structural differences, suggesting substantial species-specific regulation of RNA structure (Figure 3e). Taken together, our analyses suggest that structural changes are pervasive, reflecting that many different factors may contribute to their regulation in different circumstances.

### RNA modification and RBP binding underlie RNA structure dynamics

RNA modification and RBP-binding are important factors that are known to influence RNA structure. To disambiguate their contributions to RNA-structural dynamics, we overlaid compartment-specific RNA structuromes with RNA modifications and RBP-binding sites. Figure 4a-c shows examples of focal conformational changes around known locations of m6A modification, pseudouridylation (Ψ) and heterogeneous nuclear ribonucleoprotein C (HNRNPC) binding.

**Fig 4.**
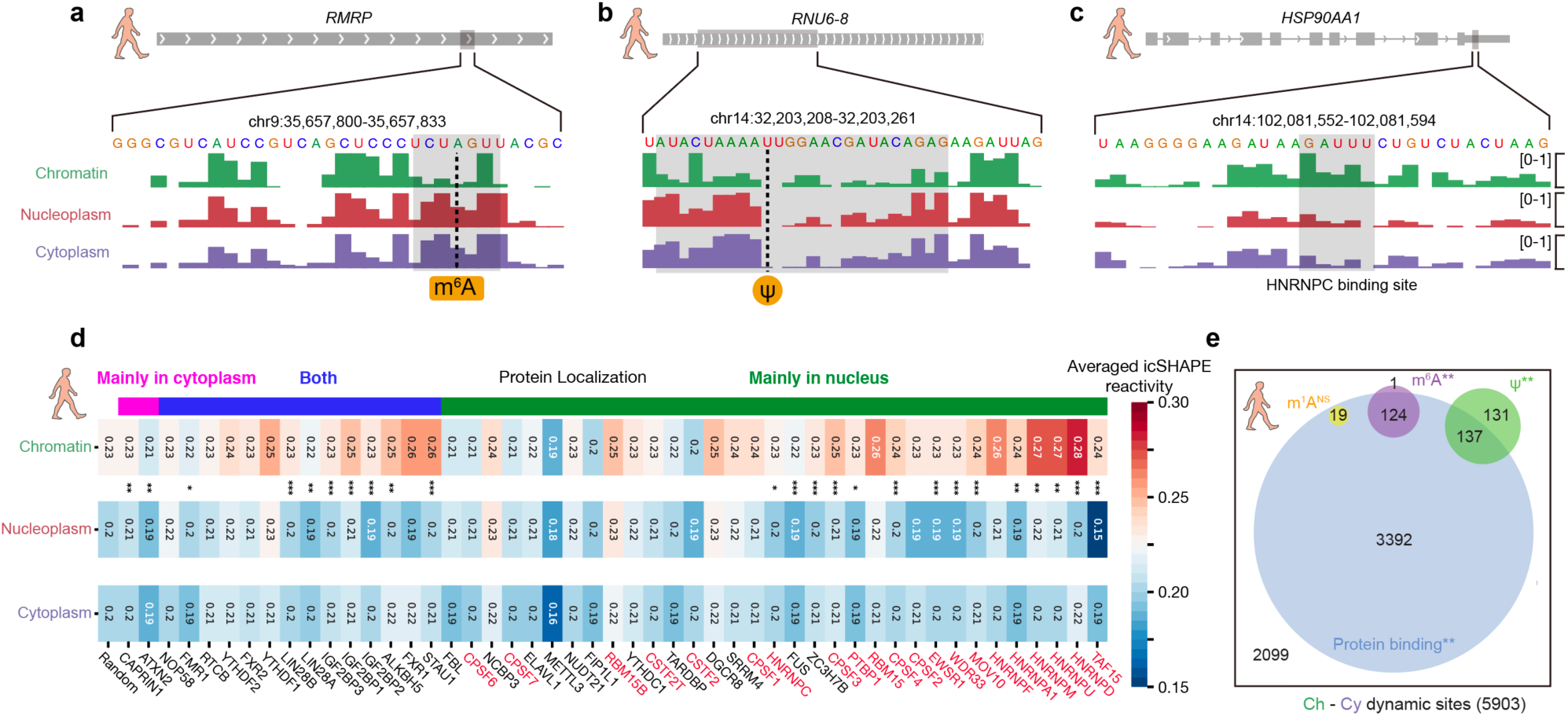
RNA modification and RBP binding underlie RNA structure dynamics. **a-c**, RNA structure dynamics at (a) an m^6^A-modified site, (b) a Ψ-modified site, and (c) an HNRNPC-binding site. Tracks show the icSHAPE score plotted along the RNA sequence. **d**, Heatmap of average icSHAPE scores in RBP binding regions in different cellular compartments, ranked by increasing structure dynamics (from left to right) between the chromatin and the nucleoplasmic fractions. Proteins are annotated by their known localizations, with chromatin-associated RBPs shown in red. P-values were calculated by single-sided Mann-Whitney U test and corrected by bonferroni method. **e**, The number and overlaps of different types of RNA modification sites and RBP binding sites in RNA structurally-dynamic regions. P-values were calculated by shuffling 1000 times. ∗ p-value < 0.05; ∗∗ p-value < 1e-3; ∗∗∗ p-value < 1e-5

As m^6^A is well known as an RNA structure switch, favoring unpairing of dsRNA^12,28^ we compared the genome-wide structures for m^6^A methylated versus non-methylated sites with the same underlying sequence motif, and confirmed similar patterns of structure destabilization in all three fractions (Figure S9a). Furthermore, the structural differences are largest in the nucleoplasm fraction, consistent with the finding that METTL3-METTL14 complex deposits m^6^A on nuclear RNA^42^. Following the structural dynamics of the same set of m^6^A sites from nucleoplasm and cytoplasm, we observed that RNA structure appears more open upon RNA migrating from the chromatin to the nucleoplasm, and thereafter remains the same (Figure 4a, S9a). Both analyses suggest that the majority of m^6^A happens in nucleus. We repeated the analysis for pseudouridylation, another abundant RNA modification generated by the isomerization of uridine, which permits hydrogen bonding to the adjacent phosphate backbone. The extra hydrogen bond can rigidify RNA structure of Ψ-modified regions^24^. We found that in general these regions have higher icSHAPE reactivity (*i.e*. less structured), suggesting that modification hinders RNA structure folding freely, which again occurs predominantly in nucleus. (Figure 4b, S9b).

All RNAs associate extensively with proteins in cells, and RNA binding protein (RBP) interactions are both sensitive to and profoundly impacts RNA structure. Taking HNRNPC as an example, we first confirmed that it bound to a stem-loop structure, inferred from more single-stranded nucleotides with flanking dsRNA (Figure 4c). We also followed the structure dynamics of the binding sites from chromatin to nucleoplasm and cytoplasm. We found that HNRNPC binding sites are more open in chromatin, also consistent with its major localization in chromatin-associated pre-RNA (Figure S9c). Our findings also suggest that HNRNPC binding could be a factor that accounts for the structural dynamics around the binding sites. Indeed, there is a significant overlap between HNRNPC binding and structurally-dynamics sites (Figure S10).

We extended the analysis to all RBPs with binding site information available from published RBP CLIP-seq experiments^43^. As shown in Figure 4d, occupancy of many RBPs are linked with RNA structural changes, while others preferentially bind to structurally-stable regions of RNA. For example, many chromatin-associated proteins (e.g. HNRNPD and others shown in red in Figure 4d) bind to more open RNA regions; these regions become more structured after dissociating from the chromatin and the proteins. In contrast, the double-stranded-binding RBP Staufen homolog 1 (STAU1), a protein that shuttles between the nucleus and the cytoplasm, appears to stabilize RNA structures upon its binding after RNA leaves chromatin. Thus, by determining the structuromes of multiple cytotopic localizations, our study provides an estimate of the relative contributions of known modification mechanisms and protein binding to RNA structure dynamics (Figure 4e). Protein binding using existing CLIP-seq data can explain most of the RNA structure dynamic sites (3392 of 5903), and many RNA modification sites with RNA structure changes overlap with protein binding sites. Our results thus suggest a complex interplay among RNA modification, protein binding and RNA structure dynamics.

### Structural analysis dissects different types of m^6^A readers

Identifying RBPs that can read RNA modifications is of fundamental significance in the study of epitranscriptomics^42^ Using our dynamic RNA structurome data to filter published CLIP-seq data, we computed the effect that m^6^A modification has on protein binding (Methods). Our analysis identified most of the known m^6^A readers, including the canonical YTH domain proteins, and the newly identified HNRNPC^28^ and the IGF2BP proteins^44^. All these readers bind to a region that contains one or more m^6^A sites stronger than a control (unmodified) site with the same m^6^A sequence motif (Figure 5a). Interestingly, the analysis also revealed several proteins with decreased bindings on modified m^6^A sites (termed “anti-readers”)^42,45^, including LIN-28 homolog A (LIN28A) and EW RNA binding protein 1 (EWSR1).

**Fig 5.**
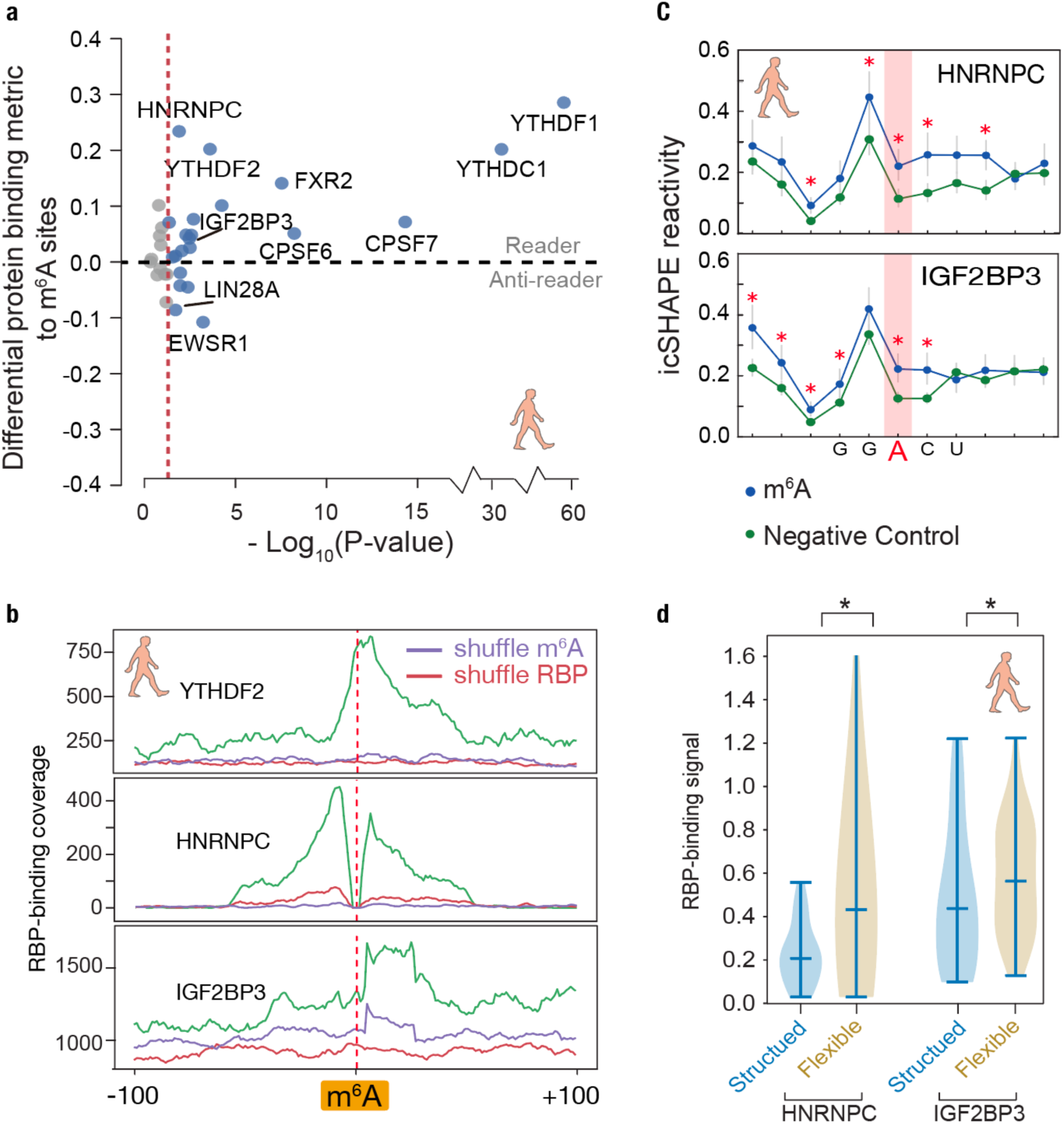
Structural analysis dissects different types of m^6^A readers. **a**, Differential RBP binding to m^6^A sites and control sites containing an m^6^A motif. P-values are calculated to show the statistical significance of the binding differences by single-ended Mann-Whiteney U test and corrected by Benjamini/Hochberg method. **b**, Metagene profiles of protein binding in m^6^A-flanking regions. **c**, Metagene profiles showing RNA structures are different between known m^6^A-modified sites and control (unmodified) sites, with the m^6^A motif around the binding sites of IGF2BP3 and HNRNPC. P-values were calculated by single-ended Mann-Whiteney U test, red stars mean p-values are less than 0.01. The error bars represent the standard error of the mean. **d**, Violin plots of RBP-binding strengths of HNRNPC and IGF2BP3 in structured and flexible regions containing a m^6^A motif. Structured and flexible regions are defined as the RBP-binding regions at the bottom 30% and top 30% of average icSHAPE scores respectively. P-values were calculated by single-ended Mann-Whiteney U test, black stars mean p-values are less than 0.05.

The precise pattern of RBP binding peaks and RNA structure at m^6^A sites can further reveal the biochemical mechanism of the m^6^A readers (Figure 5b). While the canonical readers bind most strongly directly at the m6A sites, the binding of HNRNPC and IGF2BP readers peaks at a distance. Our icSHAPE data supports a previous study that suggested that HNRNPC acts as m^6^A reader not by recognizing the N6-methyl group, but rather by binding a purine-rich motif that becomes unpaired and accessible upon nearby m^6^A modification^28^ (Figure 5c). Similarly, our RNA-structural data suggest that IGF2BP proteins (here IGF2BP3) may also read the structural changes induced by the so-called m^6^A-switch^44^ (Figure 5c). Furthermore, both HNRNPC and IGF2BP3 bind more tightly to flexible regions (Figure 5d).

To validate the “indirect reader” IGF2BP3 and the “anti-reader” LIN28A, we selected four endogenous m^6^A sites as targets. Each of the four targets contained three variants for the m^6^A site — an unmodified nucleotide, an m6A modification, and an adenosine-to-uracil mutation that mimics the disruption of base pairing (for IGF2BP3) or RBP binding (for LIN28A) (Figure 6a-b, S10a-b). We synthesized RNA constructs and used these RNA probes to retrieve RBPs from cell lysates. RNA pulldown analyses revealed that IGF2BP3 displays enhanced binding to the m6A-modified RNAs and uracil mutations relative to the unmethylated controls, confirming IGF2BP3 to be a m^6^A-switch reader (Figure 6a, S11a). Conversely LIN28A displayed reduced binding to the m6A-modified and uracil-mutant target RNAs, supporting the hypothesis that LIN28A is an anti-reader that requires an unmethylated adenosine for binding (Figure 6b, S11b).

**Fig 6.**
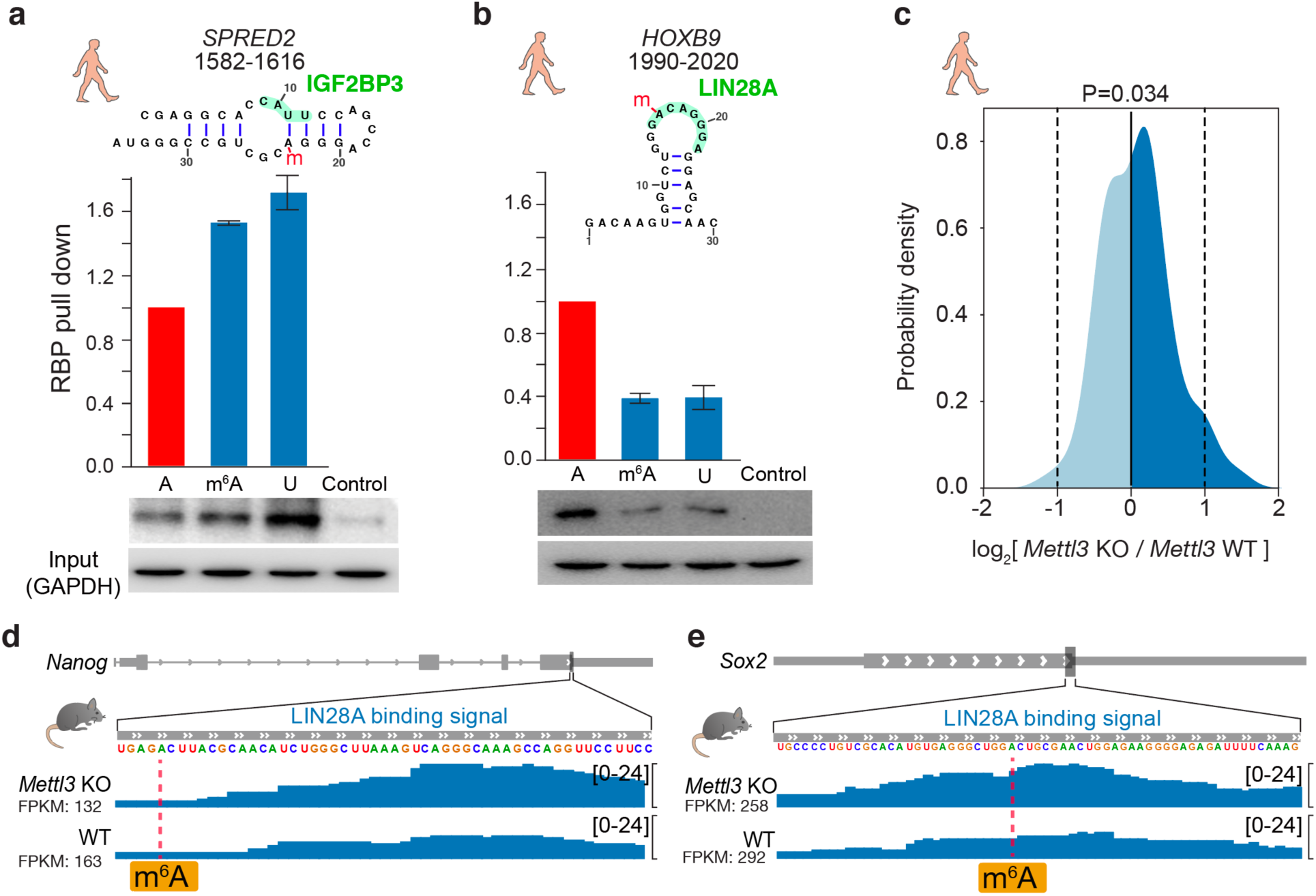
Validation of IGF2BP3 as an indirect m^6^A reader and LIN28A as an anti-reader respectively. **a-b**, RNA pull-down assays and RBP Western blots for (a) IGF2BP3 and (b) LIN28A, using RNA probes that contain unmodified A, m^6^A, and U respectively derived from the indicated positions in the transcripts. M^6^A sites are marked with a red “m”. Histograms show RNA pull-down from three replicates. Western blots are done with anti-IGF2BP3 or anti-LIN28A antibodies respectively after RNA pull down. The error bars represent standard deviation of replicates. **c.** Density plot of LIN28A binding strength (log ratio) at m^6^A sites in *Mettl3* knockout (KO) *v*ersus wild-type mES cells. P-value is calculated by single-ended t-test. **d-e**, Signal tracks of *Sox2* (sex determining region Y box 2) and *Nanog* (nanog homeobox) showing LIN28A binding at specific loci in *Mettl3* KO and wildtype mES cells.

To confirm the anti-reader role of LIN28A, we performed LIN28A CLIP-seq experiments in the wild type and the m^6^A-methyltransferase *Mettl3*-knockout mES cells^46^. Many mRNAs containing one or more known m6A site showed increased binding to LIN28A when m^6^A deposition is abrogated, relative to the negative controls (p = 0.034, t test, Figure 6c-e, S11c-d). Increased LIN28A binding is not due to increased mRNA accumulation in *Mettl3* KO ES cells (Figure 6c-e, Figure S11c-d). LIN28A is an RBP known to enforce ES cell pluripotency and suppress ES cell differentiation^47^, while m^6^A is required for stem cell differentiation^46^. The negative regulation of m^6^A on LIN28A binding is consistent with the protein’s functional roles. For example, LIN28A is a well-studied inhibitor of primary microRNA processing^48^, and m6A was recently shown to promote pri-miRNA processing^29^. Thus, the discovery of LIN28A as an m6A anti-reader potentially unifies their functional and molecular mechanisms in pluripotency, microRNA biogenesis, and post-transcriptional gene regulation.

## Discussion

Our analysis of RNA structuromes in different subcellular locations illuminated distinct RNA structural states in chromatin, nucleoplasm and cytosol. Fractionation enriched specific pools of RNAs, such as nuclear-enriched lncRNAs and pre-mRNAs including introns, thus substantially expand the scope and comprehensiveness of the RNA structuromes. Cytotopic RNA structuromes revealed the intimate connection between RNA structure and RNA processes such as transcription, translation, RNA degradation, RBP interaction and RNA modification. Through comparative analysis, we were able to dissect the role of RNA modifications and RNA-binding proteins in influencing structure, and resolved the different sets of direct and indirect RNA modification readers. We further found and validated a novel role of the pluripotency regulator LIN28A as an anti-reader for m^6^A modification.

How RNA structure is regulated *in vivo* had remained elusive, although this information is essential to revealing hidden roles of RNA structures in gene expression regulation. Our study presents the first dynamic landscape and regulation of RNA structuromes in mammalian cells. By comparative analysis we showed that the majority of the RNA structures are stable across three locations, suggesting that they have been largely determined since their biogenesis (Figure 3a-b). This structure stability could partially explain the correlations between different RNA events including transcription, translation and RNA decay (Figure 2h). Future studies involving structure perturbations that uncouple those functional correlations are required to test this hypothesis.

Nevertheless, our analysis has also revealed a large number of dynamic structural sites, which undergo conformational changes as RNAs transit from their sites of transcription on chromatin, are processed in the nucleus, and ultimately decoded in the cytoplasm. A recent study examined mRNA structure changes during zebrafish early embryogenesis and found translation to be a major driving force that shapes the dynamic mRNA structural landscape^49^. Our cytotopic data offers an opportunity to validate the finding in mammalian cells. We found that the structural dynamics between mRNAs in the chromatin fraction and the nucleoplasmic fraction is about the same as that between mRNAs in the nucleoplasmic fraction and the cytoplasmic fraction. Furthermore, the structural dynamics for mRNAs are similar to those for lncRNAs (Figure 3b). As translation remains a possible important biological process that helps to shape RNA structuromes, our observations suggest that other factors may play crucial roles that regulate RNA structure for both mRNAs and lncRNAs, in a similar fashion in mammalian cells.

Among many factors known to influence RNA structure, RNA modification and RBP-binding are important *cis*- and *trans*- regulators. Our comparative analysis illuminates their relative contributions to the observed RNA structure differences in different aspects. *In vivo* (Figure 3b) both RNA modification and RBP-binding are likely different in different compartments, whereas *in vitro* (Figure 3c) there are no RBP-binding to contribute to the structure changes. This difference in regulators may explain why RNA structures are more dynamic *in vivo*. When comparing *in vivo* to *in vitro* structure (Figure 3d) RNA modification should remain unchanged, but there are no RBP-binding to contributes to the structural differences *in vitro*, thus suggesting that RBP-binding as a whole is an important regulator of RNA structure changes. And finally, both of RBP-binding and RNA modification are likely very different in mouse and human, which may account for the big structural divergences in the two species (Figure 3e).

Finally, the specific RNA regions that undergo structural transition at each subcellular location provide direct readouts of the molecular mechanisms that shape the gene expression program. The finding of LIN28A as a m^6^A anti-reader may have implications for human disease, as both LIN28A and m^6^A have been implicated in cancer progression, germ cell development, and metabolism^50^. In the future, studying RNA structure dynamics together with RNA modifications and RBP binding in physiological states, and in the context of biological and structural perturbations, will help to elucidate the complex regulatory role of RNA structures in biology and medicine.

## Acknowledgments

We thank members of the Chang and Zhang labs for discussion. We thank Ryan Flynn for experimental advice, and Chao Dai for computational advice. This work is supported by NIH Grants No. R01-HG004361 and R35-CA209919 (to H.Y.C.), by the National Natural Science Foundation of China (Grants No. 31671355, 91740204, and 31761163007), and the National Thousand Young Talents Program of China (to Q.C.Z.). F.M.F. was supported by a NIH T32 Genomics Training Fellowship and the Arnold O. Beckman Postdoctoral Fellowship. H.Y.C. is co-founder and serves on the SAB of Epinomics and Accent Therapeutics. Some sequencing data was generated on an Illumina Hiseq 4000 that was purchased with funds from NIH (award number S10OD018220). H.Y.C. is an Investigator of the Howard Hughes Medical Institute.

## Author contributions

H.Y.C and Q.C.Z. conceived the project. L.S. and F.M.F. performed icSHAPE experiments in human and mouse cell lines respectively. P.L. and L.S. analyzed all the results. J.P.B. performed the Lin28A CLIP-seq experiments. B.L. and L.T. assisted with experiments, and W.H. assisted with analysis. Q.C.Z. and H.Y.C. supervised the project. Q.C.Z., L.S., F.M.F. and H.Y.C. wrote the manuscript with inputs from all authors.

